# A modular cranial window enabling maintainable widefield optical access in non-human primates

**DOI:** 10.64898/2026.08.02.742348

**Authors:** Wonjoon Jeong, Dongsu Kim, Jaehyun Kim, Youngjeon Lee, Raehyung Yoo, Joonyeol Lee, Myunghwan Choi, Hyoung F. Kim

## Abstract

Longitudinal optical imaging of the primate cortex requires stable yet maintainable cortical access, but existing cranial window approaches remain limited by mechanical instability and tissue responses that progressively degrade optical clarity. These challenges are amplified in primates, where large craniotomies and physiological variability complicate chronic experiments. Moving beyond conventional fixed cranial windows, we present the PRIME cranial window (Primate Reconfigurable Interchangeable Modular Enclosure), a modular platform enabling maintainable, long-term, and widefield optical access in awake macaques. Built on a modular architecture, PRIME supports non-invasive adjustment, repeated cortical access, and on-demand maintenance, including tissue removal, without surgical re-entry, while a dual-ring sealing mechanism prevents cerebrospinal fluid leakage and contamination. Combined with optimized viral delivery, implantation, and maintenance strategies, PRIME preserves cortical integrity and optical clarity for long-term optical imaging. PRIME establishes a versatile, re-accessible experimental framework for chronic, large-area optical studies of the primate brain.

## Introduction

Efforts to elucidate how the brain controls behavior have increasingly shifted from single-neuron analyses toward the study of distributed neural dynamics across space and time.^1–3^ Chronic cranial windows provide one means of repeatedly observing these dynamics across extended cortical regions and over prolonged timescales.^4–9^ However, maintaining a large optical interface over long implantation periods remains difficult owing to intracranial pressure fluctuations, tissue overgrowth, inflammatory responses, and mechanical instability.^10,11^

These challenges are particularly pronounced in primate cognitive neuroscience studies. Scientifically, higher-order cognitive and sensorimotor processes in primates often engage neural activity distributed across large cortical territories and extended timescales.^1–3,12,13^ Therefore, investigating cortical fields exceeding 1 cm over several months requires cranial windows that combine wide coverage with long-term stability.^7,14–16^ However, as window size and implantation duration increase, successful optical access depends not only on preserving initial clarity, but also on managing subsequent biological and mechanical changes at the brain–window interface.^17^

Existing cranial window designs have addressed individual aspects of this problem, but generally through trade-offs. Rigid, permanently fixed windows can provide stable attachment and optical alignment, yet offer limited access when tissue regrowth or fluid accumulation develops beneath the optical interface.^17–20^ Flexible silicone-based interfaces or smaller craniotomies can reduce mechanical loading and accommodate moderate changes in cortical position, but may restrict cortical coverage or provide limited means for restoring optical clarity once substantial tissue remodeling occurs.^5,14–16^

More recently, detachable cranial window designs have enabled window replacement and repeated cortical access.^7,21,22^ However, detachability introduces additional mechanical junctions and sealing interfaces that remain exposed to brain pulsation and persistent outward forces generated by intracranial pressure. Repeated opening and reassembly may alter contact pressure, increasing susceptibility to cerebrospinal fluid leakage, sealing failure or implant displacement, as observed in several previous systems and in our initial design. Moreover, although some detachable windows permit tissue cleaning or window replacement, they were not developed as platforms supporting standardized procedures for the systematic long-term management of tissue regrowth and pressure-related changes. Procedures for repeated cortical re-access, including tissue removal, post-implantation delivery, and neural recording, and their consequences for cortical integrity and optical performance also remain incompletely characterized. Chronic primate optical studies therefore require not merely a detachable window, but a platform that combines repeated cortical access with mechanically stable attachment, reproducible resealing, and defined procedures for long-term interface management.^17^

To address these combined requirements, we developed PRIME (Primate Reconfigurable Interchangeable Modular Enclosure), a modular cranial window platform designed to integrate the principal advantages of fixed, flexible, and detachable approaches while reducing their respective limitations. Achieving this required three integrated advances:

i. a screw-threaded modular architecture that allows repeated exchange of functional modules,
ii. a dual-ring sealing strategy that preserves a robust intracranial seal after repeated opening and resealing, and (iii) standardized maintenance procedures that enable repeated tissue removal while preserving cortical integrity and optical clarity. Rather than optimizing stability, accessibility, or sealing independently, PRIME was designed to support these functions within a single maintainable interface.

Importantly, repeated cortical re-access is central to PRIME, making procedural execution as important as device architecture. We therefore established integrated surgical and maintenance workflows and evaluated longitudinal recovery of optical access, cortical tissue responses, and post-implantation experimental reconfiguration. Together, PRIME provides a standardized yet adaptable framework for long-term, large-scale optical access to the primate cortex.

## Results

### Longitudinal optical access in non-human primates requires a maintainable framework

Longitudinal optical experiments in non-human primates (NHPs) require cortical interfaces that preserve optical access despite progressive tissue responses at the brain–window boundary. Rather than viewing chronic optical access as a one-time implantation problem, we conceived it as a recurring maintenance cycle in which tissue regrowth, fluid accumulation, and evolving experimental needs progressively impair optical access and therefore require periodic cortical re-access to restore a clear cortical view (Figure 1A). This concept distinguishes the engineering requirements for establishing initial optical access from those for repeatedly maintaining optical access to the same cortical region throughout long-term experiments.

**Figure 1.**
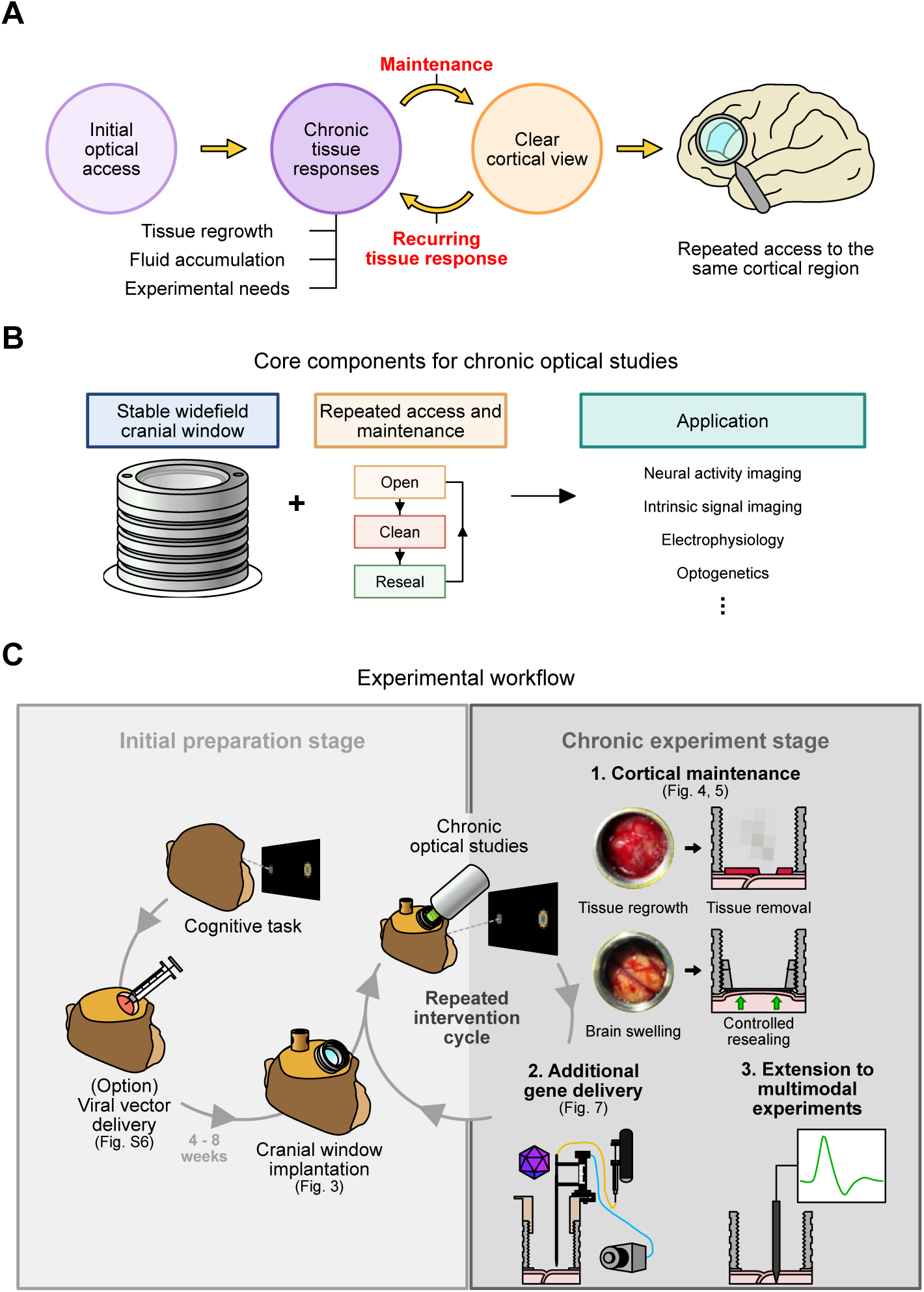
Framework for sustained cortical access during longitudinal optical studies in NHPs. (A) Framework for chronic optical window maintenance. Chronic optical access was conceptualized as a recurring cycle in which tissue responses can reduce optical clarity, and maintenance can restore a clear cortical view. (B) Core requirements for chronic NHP optical access. Stable chronic NHP optical studies require widefield optical access, repeated maintenance, and compatibility with downstream applications. (C) Workflow for chronic optical experiments. After cognitive task training and optional viral vector delivery, the cranial window was implanted for chronic optical studies. During the chronic stage, experimental needs may require repeated cortical maintenance, additional gene delivery, or extension to multimodal experiments.

To implement this concept, we established two essential components for chronic optical studies in NHPs: a stable widefield cranial window and a standardized procedure for repeated cortical re-access and maintenance, which together form a maintainable platform compatible with optical applications (Figure 1B). This platform was designed to allow the implanted interface to be repeatedly opened, serviced, and securely resealed without replacing the entire cranial implant, enabling tissue removal, viral vector delivery, and neural recording throughout long-term experiments. The system supports multiple optical modalities, including neural activity imaging, intrinsic signal imaging, electrophysiology, and optogenetic manipulation.

The experimental workflow consisted of an initial preparation stage followed by a chronic experimental stage (Figure 1C). During the initial stage, animals first completed behavioral training and, when required, underwent viral vector delivery before cranial window implantation, followed by a 4–8-week period for transgene expression. During the chronic stage, the implanted window enabled repeated cortical monitoring, tissue maintenance, additional gene delivery, and extension to multimodal optical experiments. This workflow provided the operational basis for a reconfigurable primate cranial interface that could be maintained across repeated experimental sessions.

### Design of Primate Reconfigurable Interchangeable Modular Enclosure (PRIME) cranial window

We developed the PRIME cranial window by first defining three engineering requirements for chronic optical access in NHPs: maintainable long-term optical access, widefield cortical coverage, and biological compatibility (Figure 2A). Each requirement was addressed by a corresponding design principle: a screw-threaded modular architecture enabled repeated cortical re-access and maintenance; a pressure-tolerant structure supported stable sealing across a large optical aperture; and physiologically compatible materials supported long-term contact with the cortical surface.

**Figure 2.**
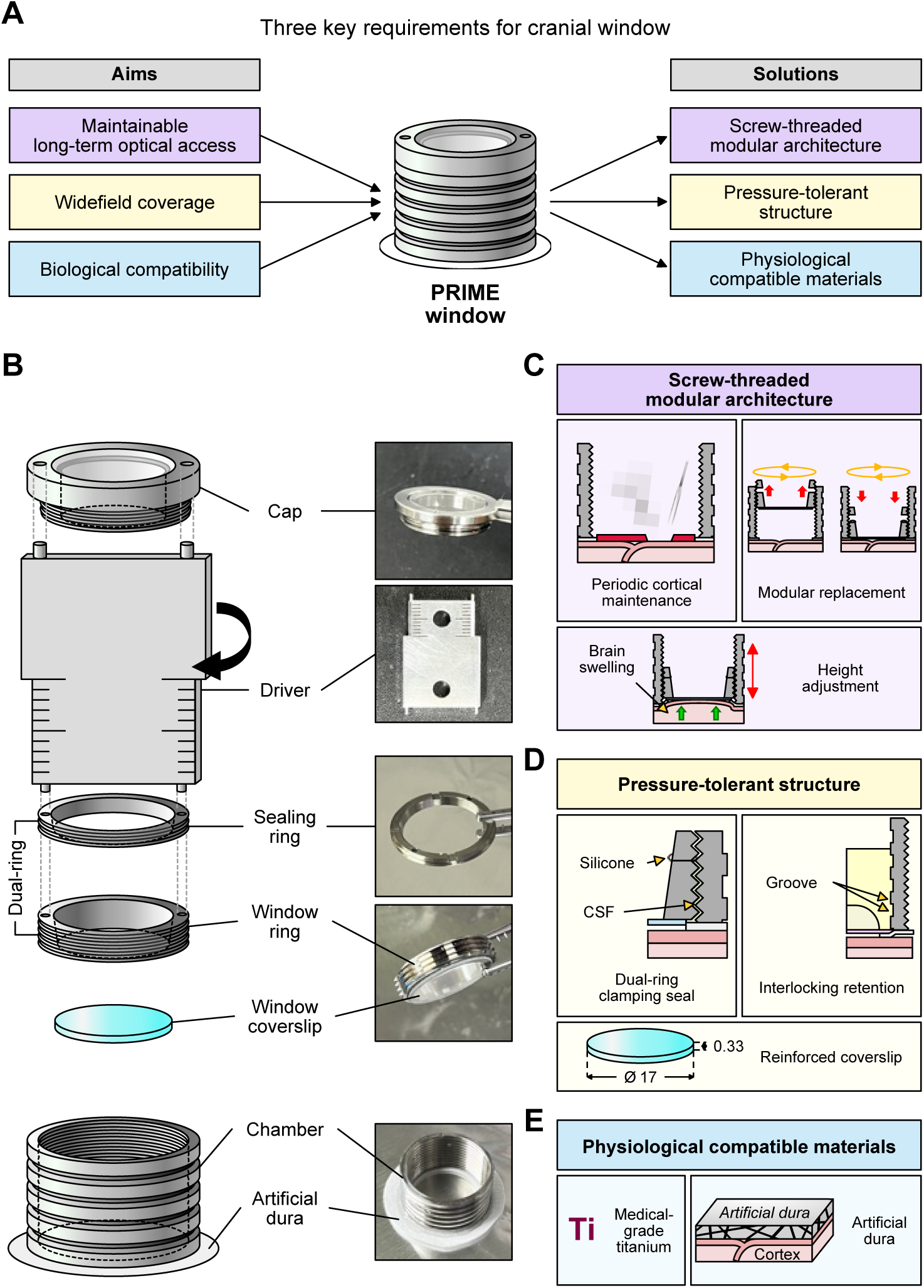
Design rationale and structural features of PRIME cranial window. (A) Design requirements and corresponding engineering solutions. The three aims were paired with corresponding solutions: a screw-threaded modular architecture for maintainable long-term optical access, a pressure-tolerant structure for stable widefield coverage, and physiologically compatible materials for biological compatibility. (B) Components of the PRIME cranial window. An exploded three-dimensional schematic and representative images show the cap, driver, sealing ring, window ring, window coverslip, chamber, and artificial dura. (C) Screw-threaded modular architecture. The threaded architecture enables periodic cortical maintenance, module replacement, and adjustment of the optical interface height in response to changes in cortical position. (D) Pressure-tolerant structure. The dual-ring clamping seal, interlocking chamber grooves, and reinforced coverslip improve sealing stability, implant retention, and structural durability. (E) Physiologically compatible materials. Medical-grade titanium and artificial dura were selected to support long-term implantation and limit direct contact between the rigid chamber and the cortical surface. All dimensions are in millimeters.

The final PRIME cranial window consists of a chamber, artificial dura, window coverslip, window ring, sealing ring, and transparent cap (Figure 2B). Its screw-threaded modular architecture allows the window ring and sealing ring to be repeatedly detached and reassembled while the chamber remains permanently fixed to the skull, enabling cortical re-access without replacement of the entire implant (Figure 2C). The threaded interface also permits adjustment of the optical window height to accommodate changes in cortical height associated with swelling, pressure variations, or cortical depression during chronic implantation.

Long-term evaluation of an initial single-ring prototype demonstrated that modularity alone was insufficient to ensure stable chronic optical access (Figure S1). Although the prototype initially provided a clear cortical view, prolonged implantation revealed cerebrospinal fluid leakage, external contamination, fibrotic tissue formation, and pressure-related mechanical failure (Figures S1D and S1E). These observations motivated further improvements in sealing stability, pressure tolerance, and structural durability.

Accordingly, the final PRIME design retained the screw-threaded modular architecture while incorporating structural features that improve sealing stability and mechanical robustness (Figure 2D). A dual-ring clamping mechanism combined with silicone sealing established a reliable fluid barrier at the chamber-window interface, preventing cerebrospinal fluid leakage and reducing the risk of contamination and infection. Grooves on the chamber exterior reinforce its coupling with dental cement, improving resistance to pressure-induced implant displacement. To enhance structural durability, the optical window was reinforced with a 0.33-mm-thick coverslip, reducing the risk of the window fracture previously observed with the thinner coverslip used in the initial prototype (Figure S1E).

Physiological compatibility was achieved using medical-grade titanium components together with an artificial dura positioned beneath the chamber (Figure 2E). The artificial dura separated the implant from the cortical surface, minimized cerebrospinal fluid intrusion, and reduced tissue ingrowth into the brain–window interface. In particular, its inner margin extended beneath the chamber opening to cover cortical regions not directly contacted by the optical window, thereby limiting fluid entry and tissue extension into the surrounding interface (Figures S2B and S2C).

Together, these engineering features integrate modular cortical access, reversible sealing, mechanical stability, and biological compatibility within a single cranial window platform. Rather than relying solely on the preservation of optical clarity after implantation, PRIME was designed to allow the brain–window interface to be reopened, maintained, and restored during chronic experiments. The complete assembly of the PRIME cranial window is shown in Figure S2.

### A controlled implantation strategy supports long-term cranial window stability

Long-term performance of a cranial window depends not only on device design but also on the implantation strategy. We therefore established a standardized surgical procedure that minimizes cortical deformation while maintaining a stable brain-window interface throughout chronic optical experiments in NHPs.

After identifying the planned sites for optical access and head fixation, the headpost was first implanted to allow behavioral training before cranial window implantation (Figures 3A and 3B). During cranial window surgery, the native dura was incised in a cruciate pattern but preserved (Figure 3C). The resulting dural flaps were temporarily reflected to expose the cortex, repositioned over the artificial dura attached to the chamber base, and sutured closed (Figures 3D and 3E). This arrangement allowed the native dura to heal over the artificial dura, while the artificial dura and chamber base acted as a physical barrier that limited the extension of regenerated dura into the brain–window interface (Figure 3E).

**Figure 3.**
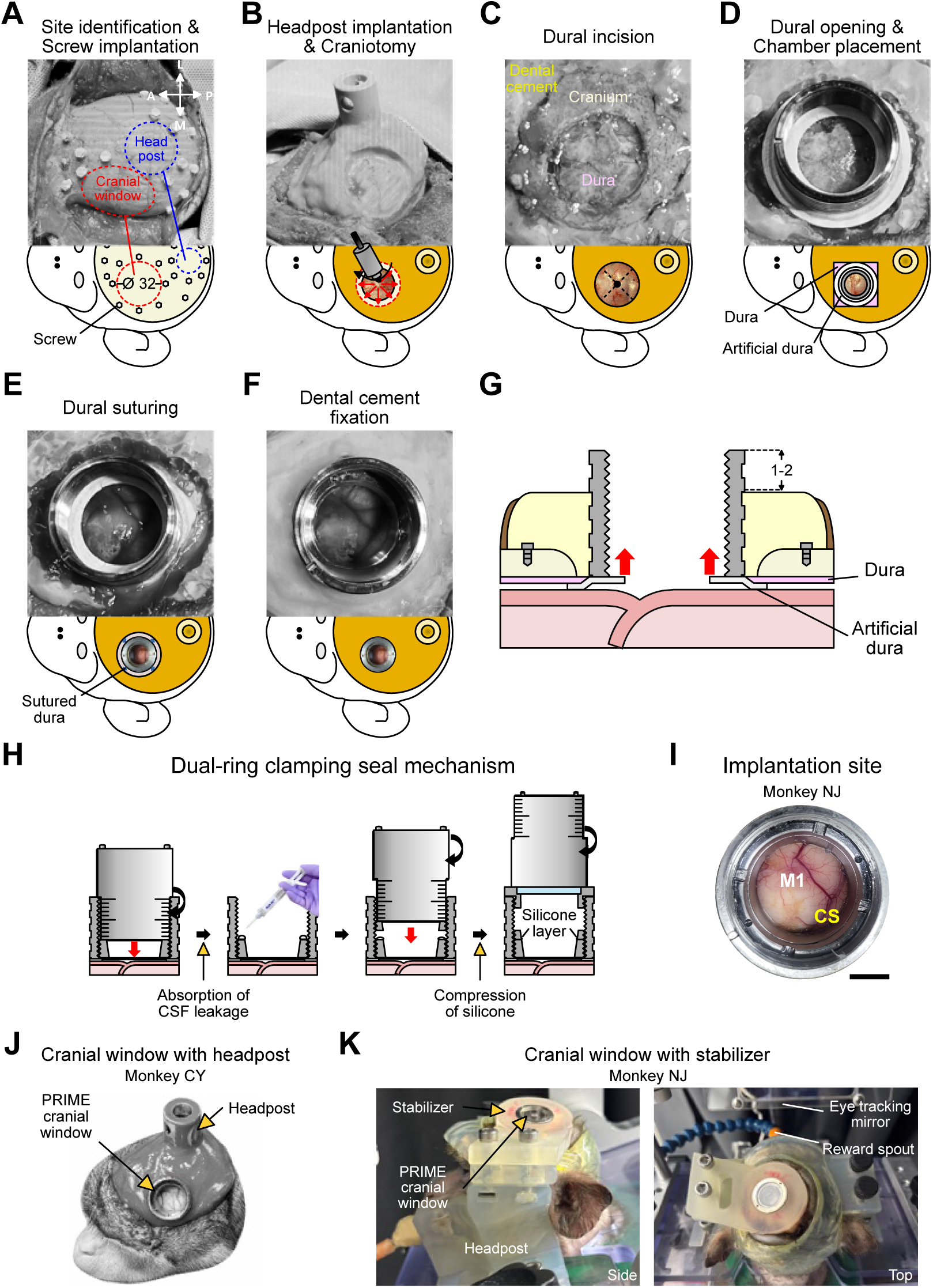
Surgical workflow for PRIME cranial window implantation. (A) Target-site planning and screw placement. MRI-based planning was used to identify the cranial window and headpost locations over the target cortical region and guide fixation-screw placement. (B) Headpost implantation and craniotomy. The headpost was implanted for behavioral training, and a craniotomy was subsequently performed at the planned cranial window site. (C) Cruciate dural incision. The native dura was incised in a cruciate pattern while preserving the resulting dural flaps for subsequent repositioning. (D) Dural opening and chamber placement. The dural flaps were temporarily reflected, and the chamber with the attached artificial dura was positioned over the exposed cortex. (E) Dural repositioning and suturing. The native dural flaps were repositioned over the artificial dura and sutured to the opposing native dural margins. (F) Dental cement fixation. The chamber was secured to the skull using dental cement. (G) Chamber positioning and stabilizer clearance. The chamber was positioned above the cortical surface to prevent direct chamber–cortex contact, and a 1–2-mm clearance was retained around the chamber when stabilizer placement was required. (H) Dual-ring clamping seal assembly. Cerebrospinal fluid released during window-ring insertion was absorbed before the sealing ring compressed the silicone layer to establish the dual-ring seal. (I) M1 implantation site. Representative optical view through the implanted PRIME cranial window over the primary motor cortex of monkey NJ. CS, central sulcus; M1, primary motor cortex. (J) Headpost-based configuration. Representative image of the PRIME cranial window and headpost in awake monkey CY. (K) Stabilizer-based configuration. Representative side and top views of the PRIME cranial window coupled to a stabilizer in monkey NJ, together with the eye-tracking mirror and reward spout. Scale bar in (I), 5 mm. All dimensions are in millimeters.

A critical implantation requirement was to avoid direct cortical compression by the rigid chamber. The chamber was therefore fixed to the skull with dental cement, leaving a 1– 2-mm clearance around the chamber and maintaining a gap above the cortical surface to prevent direct contact between the chamber wall and the cortex (Figures 3F and 3G). Direct chamber contact produced focal cortical deformation and prevented the artificial dura from conforming uniformly to the cortical surface, creating spaces into which cerebrospinal fluid, blood, or regenerated tissue could enter (Figure S3). These observations demonstrated that controlled chamber positioning is essential for maintaining a stable cranial window without deforming the underlying cortex.

After chamber fixation, the window ring was inserted and tightened to position the optical window close to the cortical surface (Figure 3H). Tightening the window ring caused cerebrospinal fluid to leak through the chamber–ring interface, relieving intracranial pressure within the sealed chamber. After pressure equilibration and cessation of leakage, Kwik-Sil silicone adhesive was applied to the window ring, and the sealing ring was fastened to compress the silicone layer. This procedure formed a secure dual-ring clamping seal across the ring–ring and chamber–ring interfaces, creating a fluid barrier while preserving the ability to repeatedly reopen and reseal the implanted interface.

Using this implantation procedure, the macaque primary motor cortex (M1) was clearly visible through the implanted optical window (Figure 3I). A transparent cap was subsequently attached to complete the cranial window assembly while allowing routine visual inspection of the implanted interface without cap removal (Figure 3H).

PRIME implantation was also compatible with alternative head-stabilization configurations. When cranial space permitted, the cranial window was implanted together with a conventional headpost for head fixation (Figure 3J). Alternatively, when cranial space was limited, a custom face-mounted stabilizer could be seated within the 1–2-mm clearance around the chamber and coupled directly to the implanted chamber to minimize head motion (Figures 3G and 3K). These complementary implantation strategies provide a flexible workflow that supports stable long-term optical access and repeated cortical re-access during awake primate experiments.

### Cortical maintenance restores optical access across phase-dependent tissue responses

After establishing stable PRIME implantation, we longitudinally monitored the cortical surface to determine whether clear optical access could be maintained over time. The cortical surface remained clearly visible for approximately 50 days after implantation, indicating that PRIME provided stable optical access during the early chronic period (Figure 4A). However, tissue regrowth emerged on day 52, progressively obscuring the cortical surface despite stable implantation. These observations demonstrate that successful implantation alone is insufficient for long-term optical access and that repeated maintenance of the brain–window interface is required.

**Figure 4.**
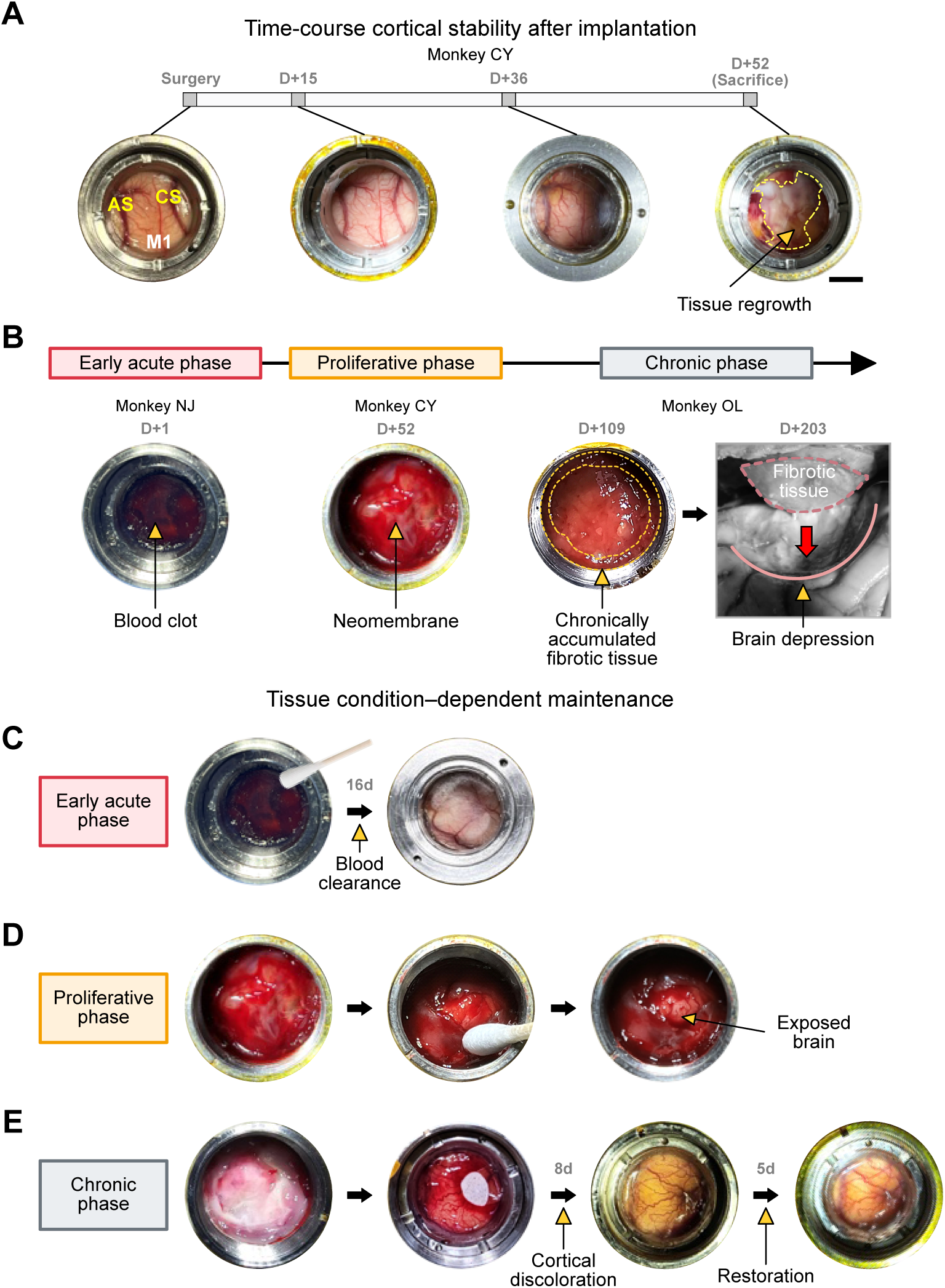
Longitudinal cortical responses and maintenance through PRIME cranial window. (A) Time course of cortical visibility after implantation. Serial observations in monkey CY show changes in cortical visibility from implantation to the development of tissue regrowth beneath the window. AS, arcuate sulcus; CS, central sulcus; M1, primary motor cortex. (B) Phase classification of post-implantation tissue responses. Tissue responses were classified according to their characteristic appearance beneath the window as early acute blood accumulation, proliferative neomembrane formation, or chronic fibrotic tissue accumulation. (C) Early acute-phase maintenance. Management of early acute blood accumulation was followed by progressive recovery of cortical visibility over the subsequent 16 days. (D) Proliferative-phase maintenance. The proliferative neomembrane was removed to re-expose the underlying cortical surface. (E) Chronic-phase maintenance. Chronic fibrotic tissue was surgically removed, followed by transient cortical discoloration and subsequent restoration of cortical visibility. Scale bar in (A), 5 mm.

Longitudinal observations from three animals revealed that tissue responses following implantation consistently progressed through acute, proliferative, and chronic phases, which were classified according to the morphological characteristics of the tissue (Figures 4B and S4). During the acute phase, residual surgical blood formed gelatinous clots on the cortical surface. In the proliferative phase, a thin neomembrane developed, partially obscuring cortical vessels while adhering weakly to the cortical surface. During the chronic phase, transparent fibrotic tissue repeatedly grew inward from the margin of the artificial dura and accumulated to form a thick fibrotic membrane. If left untreated, this tissue compressed the underlying cortex while progressively degrading optical access.

Because these phases differed in tissue composition, adhesion, and mechanical properties, maintenance procedures were tailored to each state of tissue regeneration. Acute blood clots were readily removed with a sterile cotton swab, and the cortical surface recovered its original appearance within approximately 16 days (Figure 4C). Proliferative neomembrane remained removable by gentle swabbing, restoring cortical vascular visibility (Figure 4D). In contrast, chronic fibrotic tissue formed a thick adherent membrane that required microsurgical removal using micro-scissors and micro-forceps. Removal of chronic fibrotic tissue occasionally produced focal bleeding, likely originating from angiogenic vessels within regenerated tissue adherent to the cortical surface, followed by transient yellowish cortical discoloration (Figure 4E). Nevertheless, the cortical surface recovered over time. These findings demonstrate that effective cortical maintenance requires both repeated cortical re-access and phase-specific removal strategies tailored to the mechanical properties of the regenerated tissue.

We therefore established a standardized maintenance workflow that leveraged the reconfigurable architecture of PRIME to repeatedly expose, maintain, and reseal the cortical interface without removing the skull-fixed chamber (Figure 5). Maintenance began by removing the transparent cap and disassembling the dual-ring assembly to reopen the cortical interface (Figure 5A). After cerebrospinal fluid and other accumulated fluids were cleared, regenerated tissue was removed using phase-specific maintenance procedures with appropriate tools and handling strategies. During the acute and proliferative phases, regenerated tissue was gently elevated with a vessel-dilating probe and detached by sweeping along the inner margin of the artificial dura, minimizing direct mechanical disturbance to the cortical surface (Figure 5B). In contrast, during the chronic phase, regenerated tissue had matured into a thick, adherent fibrotic membrane that required microsurgical removal. The membrane was first incised with micro-scissors and subsequently removed with micro-forceps. Residual fibrotic strands adherent to the cortical surface were carefully dissected to minimize subsequent tissue regrowth (Figure 5B).

**Figure 5.**
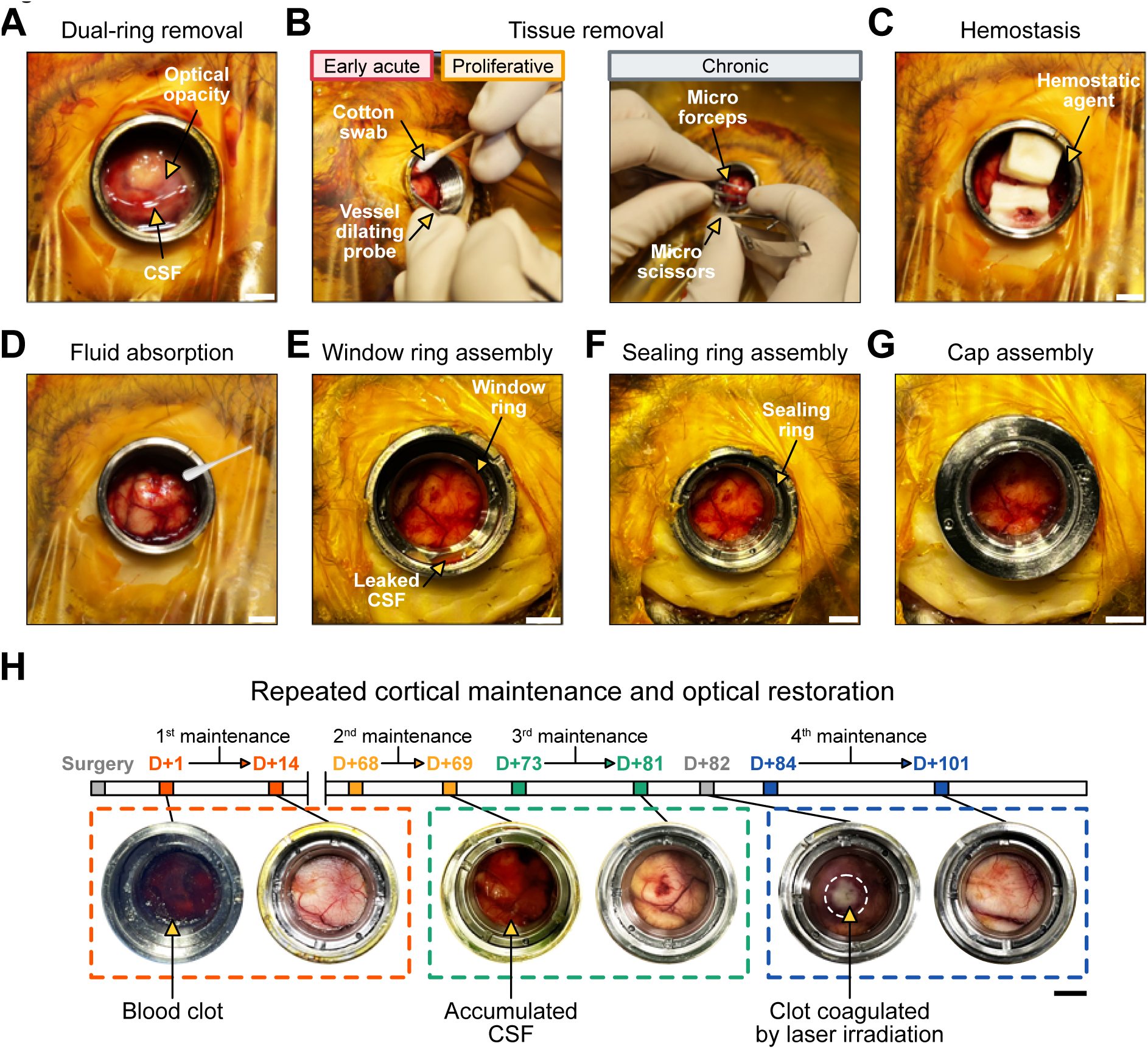
Detailed cortical maintenance procedure. (A) Dual-ring removal. Removal of the dual-ring assembly exposed optical opacity associated with cerebrospinal fluid accumulation and regenerated tissue beneath the window. (B) Phase-specific tissue removal. Early acute or proliferative tissue was removed using a cotton swab and vessel-dilating probe, whereas chronic fibrotic tissue required micro-forceps and micro-scissors. (C) Hemostasis after tissue removal. A topical hemostatic agent was applied to control bleeding after cortical maintenance. (D) Fluid absorption before reassembly. Residual fluid was absorbed before window reassembly to reduce fluid retention and bubble formation beneath the optical interface. (E) Window-ring reassembly. The window ring was reinserted, and cerebrospinal fluid released through the chamber–ring interface was repeatedly absorbed until leakage subsided. (F) Sealing-ring reassembly. Kwik-Sil was applied, and the sealing ring was tightened to re-establish the dual-ring clamping seal. (G) Final cap assembly. The cap was attached after resealing to complete cranial window reassembly. (H) Repeated maintenance and optical restoration. Longitudinal examples show recovery after blood-clot formation, cerebrospinal fluid accumulation, and localized clot coagulation following laser irradiation. Cortical visibility recovered after each maintenance intervention over the subsequent days to weeks. Scale bars, 5 mm.

After tissue removal, hemostasis was achieved,and residual fluid was cleared to leave the cortical surface slightly dry before window reassembly (Figures 5C and 5D). This preparation promoted close apposition between the optical window and the cortical surface while minimizing bubble formation during window reassembly. The window ring was then reinserted, and cerebrospinal fluid released through the chamber–ring interface during pressure equilibration and brain pulsation was repeatedly cleared until leakage subsided (Figure 5E). The dual-ring clamping seal was subsequently re-established over the membrane-cleared cortical surface, restoring a sealed brain–window interface while preserving the ability for future cortical re-access (Figure 5F). Repeated cortical re-access, phase-specific tissue removal, secure resealing, and final cap assembly enabled long-term restoration of optical access without replacing the implanted chamber (Figure 5G).

Longitudinal monitoring further showed that the maintenance workflow could be reapplied across distinct causes of optical deterioration (Figure 5H). Blood accumulation after surgery reduced optical clarity on day 1, and maintenance restored a clear cortical view by day 14. The window then remained optically clear until day 68. When cerebrospinal fluid accumulation and cortical indentation occurred after a later intervention, additional maintenance and resealing restored cortical visibility within 8 days. After recovery, photostimulation generated a localized whitish coagulated clot at a site containing residual blood products, which required another maintenance cycle; optical clarity was restored within 17 days. Across these repeated maintenance episodes, optical clarity was restored after each intervention, typically within approximately two weeks, indicating that the open–clean–reseal workflow provided a reproducible route for recovering cortical visibility without replacing the implanted chamber.

### Artificial dura biocompatibility supports chronic window performance

Longitudinal monitoring showed that regenerated tissue was largely confined beneath the window and did not broadly extend beneath the artificial dura-contacting cortex, suggesting that the artificial dura acted as a physical boundary limiting tissue ingrowth. Because the artificial dura remained in direct contact with the cortex throughout chronic implantation, we next examined whether long-term implantation induced local tissue responses by comparing GFAP and Iba1 immunoreactivity between the artificial dura-attached cortex and the contralateral intact hemisphere (Figure 6). ROIs were defined relative to the artificial dura– brain interface to assess superficial and deeper cortical responses separately (Figure S5).

**Figure 6.**
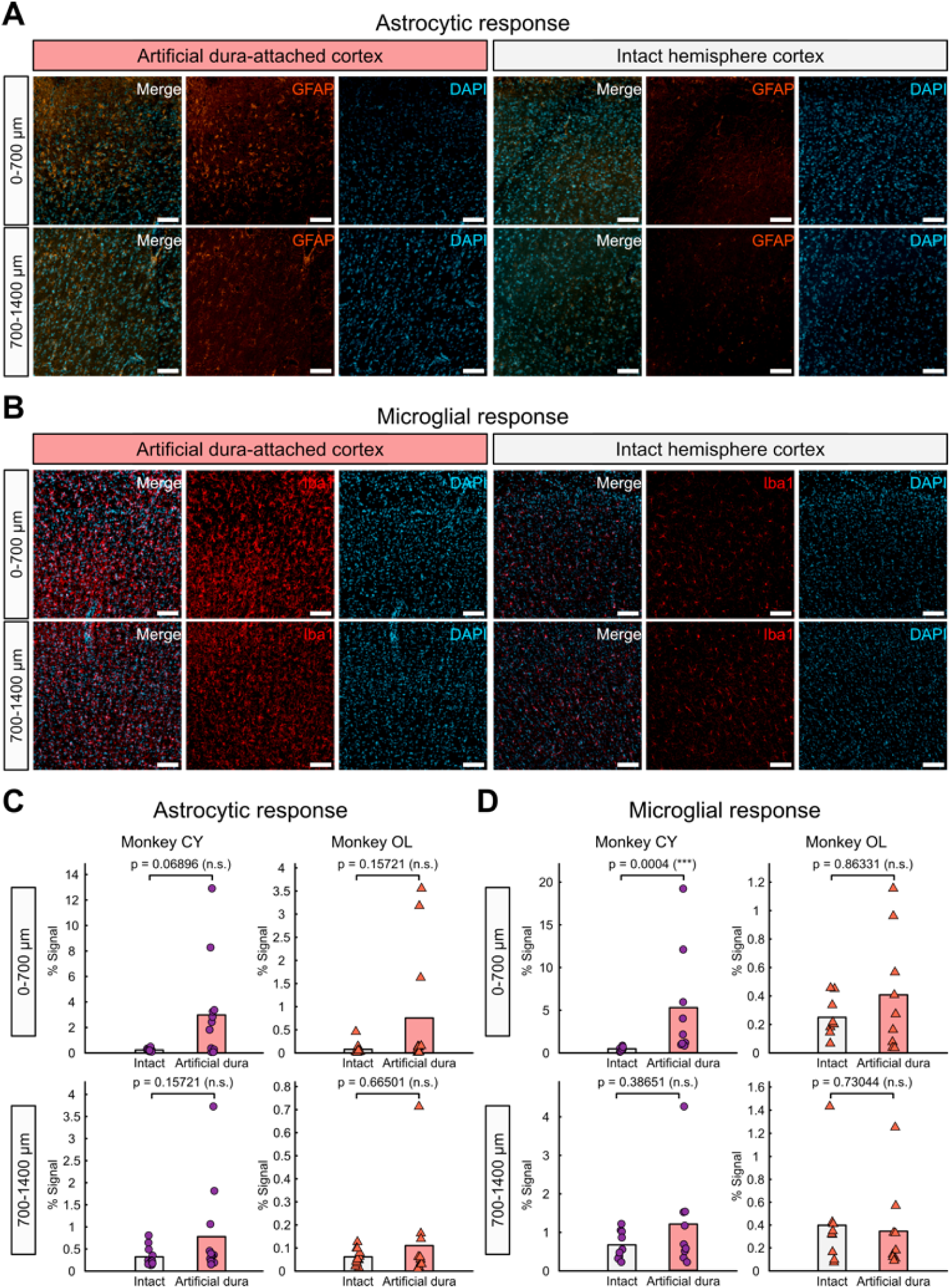
Histological assessment of cortical immune responses beneath the artificial dura interface. (A) Astrocytic response in monkey OL. Representative merged and single-channel images show GFAP and DAPI staining in the artificial dura-attached cortex and contralateral intact cortex at superficial and deeper cortical depths. (B) Microglial response in monkey CY. Representative merged and single-channel images show Iba1 and DAPI staining in the artificial dura-attached cortex and contralateral intact cortex at superficial and deeper cortical depths. (C) Quantification of astrocytic responses. GFAP-positive area fractions were quantified in 700 × 700 μm ROIs at cortical depths of 0–700 μm and 700–1,400 μm in the artificial dura-attached and intact cortical regions. (D) Quantification of microglial responses. Iba1-positive area fractions were quantified in 700 × 700 μm ROIs at cortical depths of 0–700 μm and 700–1,400 μm in the artificial dura-attached and intact cortical regions. Data were compared between artificial dura-attached and intact cortical regions using the Wilcoxon rank-sum test. ns, not significant; ***p < 0.001. Scale bars, 100 μm.

Histological analysis was performed 203 days after implantation in monkey OL and 52 days after implantation in monkey CY. GFAP immunoreactivity showed a trend toward increased astrocytic responses beneath the artificial dura, although this did not reach statistical significance (Figures 6A and 6C). Representative images from monkey OL showed no extensive GFAP-positive astrogliosis in the artificial dura-attached cortex compared with the intact hemisphere (Figure 6A). Quantification across ROIs showed no significant increase in GFAP signal in the artificial dura-attached cortex in either animal at either cortical depth (Figure 6C). Although signal intensity was frequently higher than in the contralateral control hemisphere, considerable spatial variability prevented these differences from reaching statistical significance. These results indicate that the artificial dura interface did not induce widespread astrocytic activation across the sampled cortex, although mild localized astrocytic responses may have been present.

Iba1 immunoreactivity showed a more localized pattern (Figures 6B and 6D). Representative images from monkey CY showed increased Iba1 signal in the superficial cortex beneath the artificial dura (Figure 6B). Consistent with this pattern, Iba1 immunoreactivity was significantly increased only in the superficial artificial dura-attached cortex of monkey CY (0– 700 μm, P = 0.0004, Wilcoxon rank-sum test), whereas no significant increase was detected in the deeper cortex of the same animal (Figure 6D). In monkey OL, Iba1 immunoreactivity was not significantly increased at either cortical depth (Figure 6D). Thus, although local microglial responses varied across animals following implantation, they were generally absent or limited to mild, spatially restricted activation.

Together, these histological results indicate that the artificial dura serves as an effective physical barrier while eliciting only mild and spatially restricted immune responses, supporting its suitability for long-term optical access.

### PRIME modularity enables repeated maintenance and chronic experimental reconfiguration

The preceding results demonstrated that PRIME could be repeatedly opened, maintained, and resealed while preserving chronic optical access. Because the screw-threaded modular architecture allows the optical window to be readily exchanged for other functional modules, PRIME provides a versatile platform for repeated experimental reconfiguration during chronic studies. However, chronic optical experiments may require additional gene delivery after window implantation. This may be necessary when transgene expression is insufficient or declines over time, or when additional molecular tools are needed. Gene expression may therefore be established either before or after window implantation (Figure 7A). Whereas pre-implantation delivery provides stable initial transgene expression before chronic experiments, the reconfigurable PRIME architecture enables additional genetic manipulation during the chronic experimental period. We therefore developed an interchangeable injection interface that allowed the fixed PRIME chamber to be reconfigured for post-implantation gene delivery.

**Figure 7.**
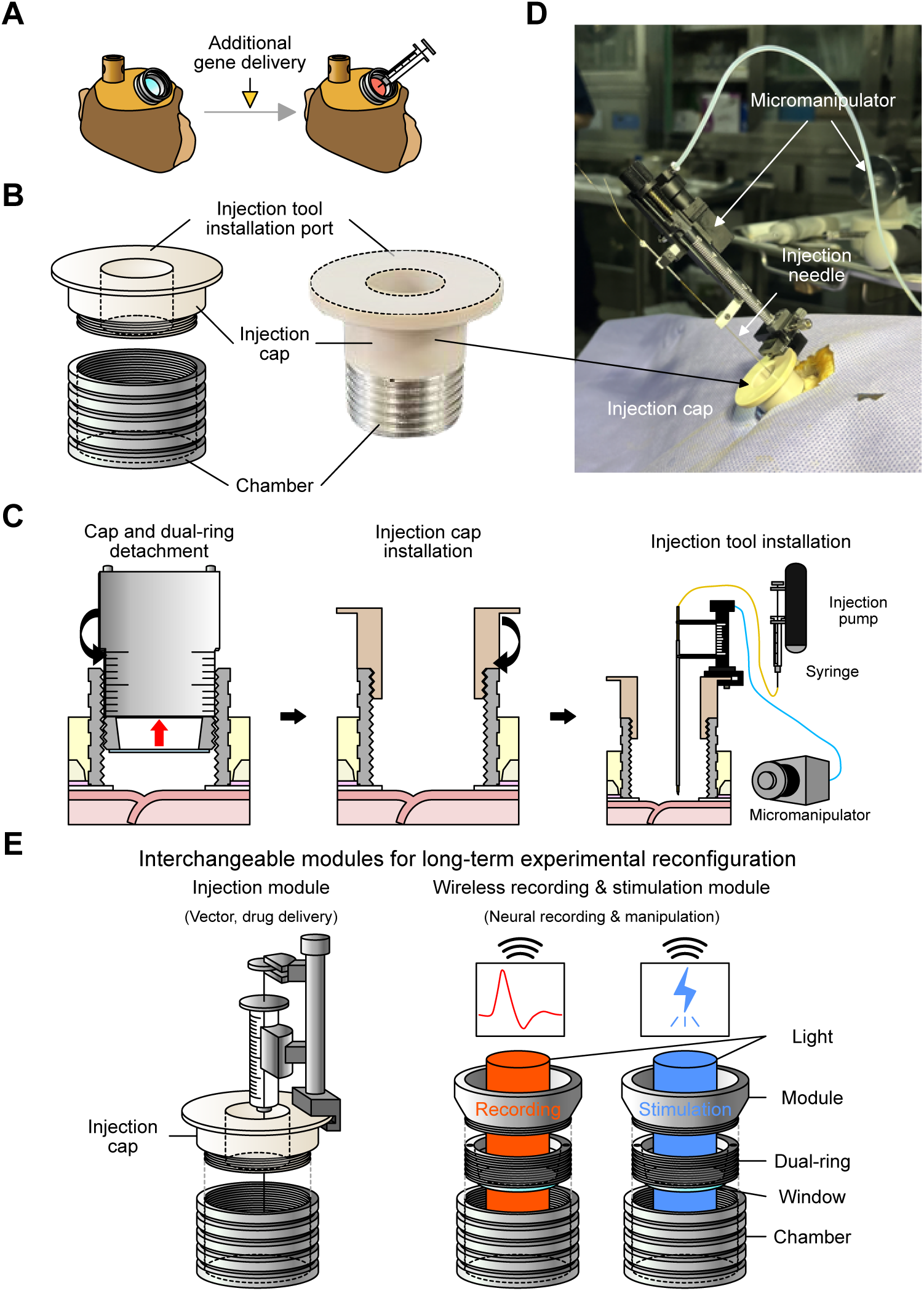
Injection module for additional gene delivery. (A) Post-implantation gene delivery. Replacement of the optical assembly with an injection interface enabled additional gene delivery through the implanted PRIME chamber. (B) Threaded injection cap. A schematic and representative image show the injection cap, injection-tool installation port, and threaded coupling to the implanted chamber. (C) Injection-module installation workflow. The cap and dual-ring assembly were removed, the injection cap was installed, and the injection tool was mounted onto the chamber. (D) Chamber-based injection setup. A representative setup shows the injection cap coupled to a micromanipulator and injection needle for controlled cortical delivery. (E) Interchangeable modules for experimental reconfiguration. The implemented injection module supports vector or drug delivery, whereas wireless recording and stimulation modules illustrate potential future extensions of the common chamber architecture.

For post-implantation gene delivery, we designed an injection cap that replaced the optical window assembly and provided a mechanically stable access port for needle insertion (Figure 7B). This configuration allowed the implanted chamber to serve as either an optical interface or an injection guide. Following removal of the optical window assembly, the injection cap was secured to the implanted chamber (Figure 7C), allowing stable alignment of the injection tool with the target site (Figure 7D). The injection needle was advanced through the chamber, and the viral vector was delivered using an injection pump. This modular configuration enabled repeated gene delivery without requiring an additional craniotomy or replacement of the implanted chamber.

In parallel, conventional pre-implantation viral delivery was used to establish initial expression in a stable intracranial environment before chronic window implantation (Figure S6). Together, the complementary pre- and post-implantation delivery strategies enable flexible establishment, monitoring, and modification of gene expression throughout chronic optical experiments in NHPs. More broadly, this capability demonstrates that PRIME is not limited to optical imaging but functions as a modular experimental platform for chronic cortical access. Beyond viral delivery, the screw-threaded architecture could be readily adapted to accommodate additional interchangeable modules, including wireless recording or stimulation devices, enabling flexible long-term experimental reconfiguration in NHPs (Figure 7E).

## Discussion

Chronic optical studies in NHPs have remained limited by challenges that cannot be overcome simply by creating a maintainable chronic cranial window. Large cortical exposure establishes a dynamic brain–window interface that undergoes continuous remodeling through tissue regrowth, cerebrospinal fluid leakage, intracranial pressure fluctuations, and mechanical stress. Consequently, long-term optical access depends not only on the initial implantation but also on whether the implanted interface can be repeatedly maintained as these changes accumulate over time. The PRIME cranial window was developed around this concept, redefining chronic windows from permanently implanted devices into reconfigurable experimental platforms that support repeated cortical access, maintenance, and long-term experimental adaptation.

### Redefining chronic windows as reproducible maintenance platforms

Previous NHP cranial windows have enabled widefield cortical imaging through large optical apertures combined with artificial dura or dura-like interfaces.^4–7,14–16^ These studies established the foundation for chronic optical interrogation of the primate cortex. Still, they were less suited for actively managing changes that emerged at the brain–window interface over time, including tissue regrowth, cerebrospinal fluid leakage, and pressure-related deformation.^6,7,14,21,22^ Rather than simply expanding optical access, PRIME addresses this limitation by integrating a screw-threaded modular architecture with standardized implantation and maintenance procedures, thereby shifting the chronic window from a static access device to a maintainable platform.

Transforming a passive access device into an actively maintainable interface required PRIME to satisfy two coupled requirements: reversible cortical access and reliable isolation of the intracranial environment. The screw-threaded architecture enabled the optical interface to be reconfigured without disturbing the skull-fixed chamber, but this modularity also introduced component boundaries that could become vulnerable to leakage. Our experience with the initial single-ring configuration therefore suggested that silicone applied passively around a fixed interface may not provide sufficiently stable sealing for a system intended for repeated access (Figure S1).^23^ The dual-ring design addressed this trade-off by converting Kwik-Sil into a removable compression seal rather than a permanently applied sealant, allowing cortical accessibility and environmental isolation to function as complementary design features.

Mechanical design alone was insufficient for stable long-term access. Consistent implantation and maintenance were also required, as procedural variation could affect window–cortex positioning, fluid control, and resealing quality.^21–23^ PRIME therefore integrates the hardware with a defined workflow intended to reduce procedural variability. The longitudinal observations provided a practical test of whether this integrated device-and- workflow approach could restore optical access after deterioration of the brain–window interface. Repeated maintenance returned deteriorated interfaces to an imageable state, with optical clarity recovering within approximately one to two weeks across the maintenance events examined here. These findings indicate that deterioration of cortical visibility need not define the experimental endpoint. PRIME instead provides a controlled route for restoring optical access, while its defined operating workflow is intended to support consistent implementation across animals and facilitate future adoption by other laboratories.

### Scaling chronic optical interfaces across cortical curvature and window size

A key consideration for scaling chronic cranial windows is the geometric mismatch between a flat optical interface and the curved cortical surface.^24,25^ In the present design, the artificial dura extended inward to form an annulus beneath the non-optical titanium portion of the window ring, thereby preventing direct contact between the cortex and the titanium surface surrounding the optical aperture (Figure S2). Cerebrospinal fluid, blood products, or regenerated tissue may accumulate within this space and compromise the surrounding brain– window interface.^26^ To address this issue, the artificial dura was designed with a 15-mm inner diameter, slightly larger than the optical aperture, allowing its inner margin to extend beneath the window ring and cover cortical regions not directly contacted by the optical window. This inward extension limited cerebrospinal fluid entry and the encroachment of regenerated tissue into the hidden space beneath the ring, thereby helping to delay tissue regrowth around the imaging interface.

Close window–cortex apposition within the imaging aperture was also important for delaying tissue regrowth.^21^ Persistent separation can provide space for fluid accumulation and more rapid tissue remodeling, whereas excessive contact pressure can produce cortical indentation or deformation. The present design therefore relied on adjustable window positioning to maintain close apposition while limiting excessive cortical compression. This balance will become increasingly important as the optical aperture is enlarged, since a wider flat window will encounter greater variation in cortical curvature and may create larger regions of incomplete contact.^27^ Future designs may therefore require curved or conformable optical interfaces, region-specific window geometries or more spatially controlled adjustment of window–cortex contact.^24–26^ Addressing these factors will be necessary to expand cortical coverage while preserving optical clarity and minimizing tissue disturbance over long implantation periods.

### Biological compatibility of the artificial dura-contacting cortex

The mechanical role of the artificial dura must be accompanied by preservation of the biological state of the cortex that remains in long-term contact with it. Chronic window performance therefore cannot be judged solely by optical clarity or successful restoration after maintenance. The artificial dura-contacting cortex must also remain suitable for physiological interpretation, as astrocytic and microglial responses can alter neuronal excitability, synaptic function, vascular homeostasis, and neuronal survival.^28–33^

The absence of broad glial activation, together with the restriction of the observed microglial response to the superficial cortex, suggests that the artificial dura-associated response remained spatially localized rather than extending broadly into the underlying tissue. Nevertheless, this pattern should not be interpreted as an absence of glial responses. A significant superficial Iba1 increase was observed in one animal, and numerically higher GFAP- or Iba1-positive area fractions were present in some individual ROIs even when the corresponding comparisons were not statistically significant. The artificial dura interface should therefore not be regarded as biologically inert. Moreover, GFAP- or Iba1-positive area fractions may remain relatively unchanged despite alterations in cellular activation state. Astrocytic hypertrophy or scar-like organization and microglial soma enlargement or reduced process ramification could reveal interface-associated responses not captured by area-based quantification alone.^29,34,35^

Even localized glial responses could modify the cortical microenvironment through changes in inflammatory signaling, vascular regulation, blood–brain barrier integrity, or synaptic support, potentially affecting neuronal physiology and the interpretation of optical signals acquired through the window.^31,36,37^ Biological compatibility should therefore be evaluated through complementary measures of glial morphology, neuronal and vascular integrity, blood–brain barrier status, and inflammatory signaling, rather than through GFAP and Iba1 area fractions alone.^38^ As the present histological analysis was cross-sectional, it cannot distinguish a stable interface adaptation from a transient maintenance-associated response or progressive inflammation. Longitudinal in vivo assessment followed by terminal histological analysis would help resolve these possibilities and determine whether localized responses remain stable during extended optical experiments.

### Extending PRIME beyond optical access and future implications

Beyond optical maintenance, PRIME allowed the function of the implanted interface to be changed after implantation. Replacing the optical module with an injection cap converted the chamber into an interface for post-implantation viral vector delivery. This implementation provides proof of principle that a common chamber architecture can support different experimental modules at successive stages of a study. Future modules could, in principle, support wireless electrophysiological recording, photostimulation, local drug delivery, or combined optical–electrical approaches, although these applications remain to be experimentally validated.^15,22,39–42^ Such modules could also combine cortical optical imaging with depth recording or stimulation of subcortical structures, including the basal ganglia. This configuration could facilitate investigation of how cortical population dynamics interact with striatal computations during value-guided, flexible, and habitual behaviors, although this application remains prospective.^43–48^

This flexibility is particularly relevant to longitudinal NHP studies, in which experimental requirements may evolve over time.^22,40,49^ Rather than defining all experimental capabilities at the time of implantation, an interchangeable module system could allow the interface to be adapted as the requirements of a longitudinal study evolve. This approach may enable multiple experimental modalities to be flexibly deployed within the same implanted preparation, reducing the need to predetermine the complete experimental configuration before implantation.

Although PRIME was developed for experimental use in NHPs, the broader principle of a chronically implanted interface capable of accepting replaceable functional modules may also be relevant to future human brain interfaces. Potential applications could include repeated cortical monitoring, localized therapeutic delivery, electrophysiological recording, or stimulation.^50–52^ Direct clinical translation would nevertheless require substantial redesign to meet human anatomical and neurosurgical constraints, together with long-term evaluation of infection risk, mechanical stability, tissue compatibility, and clinical safety. Thus, the translational relevance of PRIME lies primarily in its reconfigurable interface concept rather than in the direct application of the present device to humans.

### Methods Animal models

All experimental and animal care procedures were approved by the Seoul National University Institutional Animal Care and Use Committee (Approval Nos. SNU-240906-6-2 and SNU-250911-1). Experiments were conducted in one male rhesus macaque (*Macaca mulatta*), OL, weighing 13.5 kg, and two female cynomolgus macaques (*Macaca fascicularis*), CY and NJ, weighing 5.5 and 4.5 kg, respectively. Animals were housed individually under controlled environmental conditions with ad libitum access to food and water.

### Fabrication and assembly of the PRIME cranial window

All PRIME cranial window components were custom-fabricated from medical-grade titanium. The optical interface and artificial dura-attached chamber were assembled separately before final cranial window assembly (Figure S2).

### Optical interface assembly

The optical interface consisted of a window ring bonded to a circular Gorilla Glass coverslip (Corning Inc., Corning, NY, USA), which was custom-processed by SXET Glass (Shenzhen, China) (Figure S2A). A thin layer of silicone adhesive (KN-300X, Shenzhen Kanglibang Technology Co., Ltd., Shenzhen, China) was applied to the underside of the window ring, and the coverslip was aligned to the ring during bonding. The assembly was placed coverslip-down and compressed with a weighted load to provide uniform axial pressure during curing. The bonded optical interface was cured at room temperature for 24 hr. After curing, excess silicone extending beyond the coverslip boundary was removed with a blade, and residual debris was cleared with compressed air.

### Artificial dura attachment

The chamber was prepared by attaching an artificial dura (Neuro-Patch, Aesculap AG, Tuttlingen, Germany) to its underside (Figure S2B). The artificial dura was shaped into an annular structure using a custom punch with an inner diameter of 15 mm and an outer diameter of 27 mm. Silicone adhesive was applied to the bottom surface of the chamber, and the artificial dura was aligned and bonded to the chamber. Additional adhesive was applied along the interface boundary to reinforce the seal. The construct was compressed with a weighted load and cured at room temperature for 24 hr. After curing, excess adhesive protruding into the chamber interior was removed to prevent interference with subsequent vertical adjustment of the window ring. The artificial dura was then radially incised in eight directions using surgical scissors to allow smooth movement of the window ring during assembly. The final artificial dura-attached chamber is shown in Figure S2C.

### Surgical implantation of the PRIME cranial window

Animals were prepared for cranial window implantation under aseptic surgical conditions. Dexamethasone (1 mg kg^-1^, i.m.) was administered 12 h before surgery. General anesthesia was induced with ketamine (5–10 mg kg^-1^, i.m.) and medetomidine (Domitor; 0.04– 0.08 mg kg^-1^, i.m.) and maintained with inhaled isoflurane throughout the procedure. Ceftriaxone (typically 35 mg kg^-1^, i.v.) and acetaminophen (5–10 mg kg^-1^, i.v.) were administered intraoperatively for antibiotic prophylaxis and analgesia, respectively.

The planned cranial window and head-fixation sites were determined based on the target cortical region. In animals that underwent behavioral experiments before window implantation, the headpost was implanted first to permit head fixation during task training. At the time of PRIME implantation, dental cement covering the planned cranial window site was removed as needed, and the target cortex was exposed by craniotomy.

After craniotomy, the native dura was incised in a cruciate pattern but was not permanently removed. Mannitol (0.5–1.0 g kg^-1^, i.v.) was administered when intraoperative brain swelling was observed. The dural flaps were temporarily reflected to expose the cortical surface. Hemostasis was achieved using a topical hemostatic agent (Vetiblock, EMVENTER LLC, Oakland Gardens, NY, USA), and the cortical surface was kept moist with sterile saline throughout the procedure. The chamber with the attached artificial dura was then positioned over the target region. The native dural flaps were repositioned over the artificial dura and sutured to the corresponding native dural margins. The chamber was positioned slightly above the cortical surface to prevent direct compression of the brain by the rigid chamber wall. Dental cement was then applied to secure the chamber to the skull while preserving the intended circumferential clearance when a window-matched stabilizer was required.

After chamber fixation, the window ring was inserted and adjusted to position the optical interface close to the cortical surface. Cerebrospinal fluid leaking through the chamber– ring interface during tightening was repeatedly absorbed until the leakage subsided. Kwik-Sil silicone adhesive (World Precision Instruments, Sarasota, FL, USA) was then applied to the upper surface of the window ring. The sealing ring was tightened to compress the silicone and establish a dual-ring clamping seal. Finally, the cap was attached to complete the cranial window assembly. The implanted window was visually inspected to confirm cortical visibility, sealing integrity, and the absence of overt chamber-induced cortical compression.

After surgery, ceftriaxone (35 mg kg^-1^, i.m.) was administered once daily for 1 week. Acetaminophen (5–10 mg kg^-1^) was administered orally for 3 days for postoperative analgesia.

### Cranial window inspection and maintenance

The cranial window was inspected periodically under anesthesia or in awake head-fixed conditions to assess optical clarity, fluid accumulation, and tissue regrowth. Maintenance was performed when optical access was reduced by accumulated blood products, fluid, or regenerated tissue within the window cavity.

Before maintenance, the cap, chamber exterior, and surrounding skin were disinfected with betadine followed by 70% ethanol. After cap removal, the chamber interior was disinfected in the same manner. If inspection indicated that cortical re-access was needed because of blood or fluid accumulation, optical opacity, or tissue regrowth beneath the window, the sealing ring was unscrewed, and residual Kwik-Sil was removed using sterile forceps. The window ring was then removed gradually to expose the cortical surface while minimizing abrupt pressure changes.

After reopening the cortical interface, accumulated cerebrospinal fluid, blood products, and other fluids were absorbed using sterile absorbent material. Regenerated tissue was removed using tools selected according to tissue consistency and adhesion to the cortical surface (Figures 4C and 5). Loosely accumulated blood products and early proliferative tissue were removed with minimal mechanical force using sterile cotton swabs or a vessel-dilating probe. Tissue attached along the inner margin of the artificial dura was gently lifted and detached along the artificial dura boundary to minimize direct disturbance to the cortical surface. Thick or adherent fibrotic tissue was incised with micro-scissors and removed with micro-forceps. Tissue strands connected to the cortical surface were carefully dissected to reduce residual tissue that could serve as a scaffold for subsequent regrowth.

The cortical surface was irrigated with sterile saline throughout the procedure. When bleeding occurred during tissue removal, hemostasis was achieved using a topical hemostatic agent. Residual fluid was then absorbed to create a clear and slightly dry cortical surface before reassembly, reducing trapped fluid or air bubbles between the optical window and the cortex.

After maintenance, residual fluid was absorbed, and the optical interface was reassembled. The dual-ring clamping seal was re-established, as described for the implantation procedure, and the cap was replaced. Newly sterilized components were used when replacement was required, and animals received postoperative anti-inflammatory and antibiotic treatments as required.

### Viral vector delivery

Viral vectors were delivered into the primary motor cortex (M1) either before cranial window implantation (Figure S6) or through the implanted PRIME chamber (Figure 7), depending on the experimental stage. Injections were performed using custom stepped injection needles or Hamilton syringes. After vector delivery at each site, the needle was left in place for 5–10 min before being withdrawn or moved to the next site to allow tissue absorption and reduce backflow.

### Pre-implantation viral vector delivery

Pre-implantation viral vector delivery was performed in monkey NJ before PRIME cranial window implantation (Figure S6). This approach was used to allow viral expression before chronic window placement and to avoid additional cortical manipulation after tissue remodeling had occurred. All procedures were performed under general anesthesia and aseptic conditions. The target cortical region was identified using MRI guidance and the planned future cranial window field of view.

After scalp incision and skull exposure, the planned cranial window boundary and injection region were marked on the cranium. Anchoring screws were placed around the planned implantation site to support subsequent cranial window fixation. A craniotomy was performed over the target cortical region using an ultrasonic surgical device (Surgystar X, Dmetec Co., Ltd., Bucheon, Republic of Korea), and the bone flap was preserved for later repositioning. After temporary removal of the bone flap, a cruciate dural incision was made to expose the cortical surface. The cortical vasculature was monitored under a surgical microscope during cannula insertion to avoid vascular damage, and the cortical surface was kept moist with sterile saline throughout the procedure.

For stereotaxic injection, a Hamilton syringe was coupled to a microinjection pump (UMP3, World Precision Instruments, Sarasota, FL, USA) and mounted on a stereotaxic frame (Figure S6G). A stainless-steel sleeve was attached around the injection needle, leaving the distal 1 mm of the needle tip exposed to form a stepped structure (Figure S6H). The injection needle had an inner diameter of 130 μm and an outer diameter of 470 μm, and the sleeve had an inner diameter of 520 μm and an outer diameter of 780 μm. The cortical vasculature was monitored under a surgical microscope during cannula insertion to avoid vascular damage, and the cortical surface was kept moist with sterile saline throughout the procedure.

After viral vector injection, the dura mater was returned to its original anatomical position and sutured. The preserved bone flap was repositioned, and bone powder (OSTEON III, Dentium Co., Ltd., Suwon, Republic of Korea) was packed into the surrounding craniotomy gap to stabilize the site. The surgical region was then sealed using dental cement (UNIFAST Trad, GC Corporation, Tokyo, Japan). Following recovery from the injection surgery, PRIME cranial window implantation was performed after a 4–8-week expression period.

### Post-implantation injection through the PRIME chamber

Post-implantation viral vector injection was performed in monkey OL after PRIME cranial window implantation (Figure 7). Before injection, the cap, sealing ring, and window ring were sequentially removed from the implanted chamber, and a threaded injection cap was attached to the chamber. A micromanipulator (MO-974A, Narishige, Tokyo, Japan) was mounted onto the injection cap to control insertion of the injection needle through the chamber.

For chamber-based injection, viral vectors were delivered using a custom stepped injection needle. A polyimide sleeve was attached around the silica injection needle, leaving the distal 1 mm of the needle tip exposed to form the stepped structure. The silica injection needle had an inner diameter of 130 μm and an outer diameter of 470 μm, whereas the polyimide sleeve had an inner diameter of 830 μm and an outer diameter of 1.27 mm.

### Immunohistochemistry

Monkeys OL and CY were deeply anesthetized with sodium pentobarbital (80 mg kg^−1^, i.v.) and transcardially perfused with 0.01 M phosphate-buffered saline (PBS; pH 7.4), followed by 4% paraformaldehyde (PFA) in PBS. Brains were extracted and post-fixed in 4% PFA at 4 °C for 7 days. For cryoprotection, brains were sequentially immersed overnight at 4 °C in PBS containing 0.1% sodium azide supplemented with 10% and 20% glycerol. Coronal brain blocks were sectioned at a thickness of 40 μm using a sliding microtome (SM2010R; Leica Biosystems, Nussloch, Germany), and sections were mounted onto glass slides before immunostaining.

To assess neuroinflammatory responses, microglia and astrocytes were labeled using antibodies against ionized calcium-binding adaptor molecule 1 (Iba1) and glial fibrillary acidic protein (GFAP), respectively. Mounted sections were permeabilized and blocked in PBS containing 0.3% Triton X-100 and 10% normal goat serum (01-6201; Invitrogen, Carlsbad, CA, USA) for 30–60 min at room temperature. Sections were then incubated overnight at 4 °C with primary antibodies diluted in PBS containing 0.3% Triton X-100 and 2% normal goat serum: mouse anti-Iba1 (1:500; MA5-27726; Thermo Fisher Scientific, Waltham, MA, USA) and rabbit anti-GFAP (1:300; ab68428; Abcam, Cambridge, UK).

After primary antibody incubation, sections were washed three times in PBS for 5–10 min per wash and incubated for 1–2 hr at room temperature with secondary antibodies diluted in PBS containing 0.1% Triton X-100 and 2% normal goat serum: goat anti-mouse IgG (H+L) Alexa Fluor 647 (1:500; A-21235; Thermo Fisher Scientific) and goat anti-rabbit IgG (H+L) Alexa Fluor 568 (1:500; A-11036; Thermo Fisher Scientific). Sections were then washed three times in PBS and coverslipped using VECTASHIELD HardSet mounting medium with DAPI (H-1500; Vector Laboratories, Newark, CA, USA).

### Image acquisition and analysis

Fluorescence images were acquired using a slide-scanning microscope (Axioscan 7; Carl Zeiss AG, Oberkochen, Germany). Images were imported into ZEISS ZEN 3.13 software (Carl Zeiss Microscopy GmbH, Jena, Germany) for visualization and region selection. To evaluate the spatial distribution of astrocytic and microglial responses across cortical depths, square regions of interest (ROIs; 700 × 700 μm) were defined in each cortical section. To minimize optical artifacts and non-specific background signals at the outermost tissue boundary, the upper edge of the superficial ROI was positioned immediately below the pial surface. The superficial cortical region was defined as 0–700 μm from the pial surface, and the deeper cortical region was defined as 700–1,400 μm by placing a second ROI directly beneath the superficial ROI along the cortical depth axis (Figure S5).

Quantification of immunoreactive area fractions was performed using ImageJ. For each ROI, GFAP- and Iba1-positive areas were segmented using a standardized thresholding procedure applied consistently within each staining channel and animal. The percentage area fraction of GFAP or Iba1 signal was calculated for each ROI and used for statistical comparison between the artificial dura-contacting cortex and the contralateral intact cortex.

### Statistics

Statistical comparisons were performed using the Wilcoxon rank-sum test. For each marker, comparisons were performed separately for superficial (0–700 μm) and deeper (700–1,400 μm) cortical regions within each animal. All statistical analyses were conducted in MATLAB, and statistical significance was defined as P < 0.05.

## Acknowledgements

This work was supported by the Bio&Medical Technology Development program (RS-2025-02263832), the Basic Science Research Program (RS-2024-00339355), the Global-LAMP program (RS-2023-00301976), and the ASTRA program (RS-2024-00436783) through the National Research Foundation (NRF) of Korea. We thank D.I. Ko for technical support and members in Institute of Laboratory Animal Resources (ILAR), SNU for technical assistance.

## Author Contributions

H.F.K. supervised the entire project. W.J., D.K., and H.F.K. conceptualized and designed the experimental setups. J.K., Y.L., and M.C. participated in gene delivery study. W.J., D.K., R.Y., J.L., and H.F.K. performed the surgery. W.J., D.K., and H.F.K wrote the first draft and the final manuscript.

---

**Figure S1.**
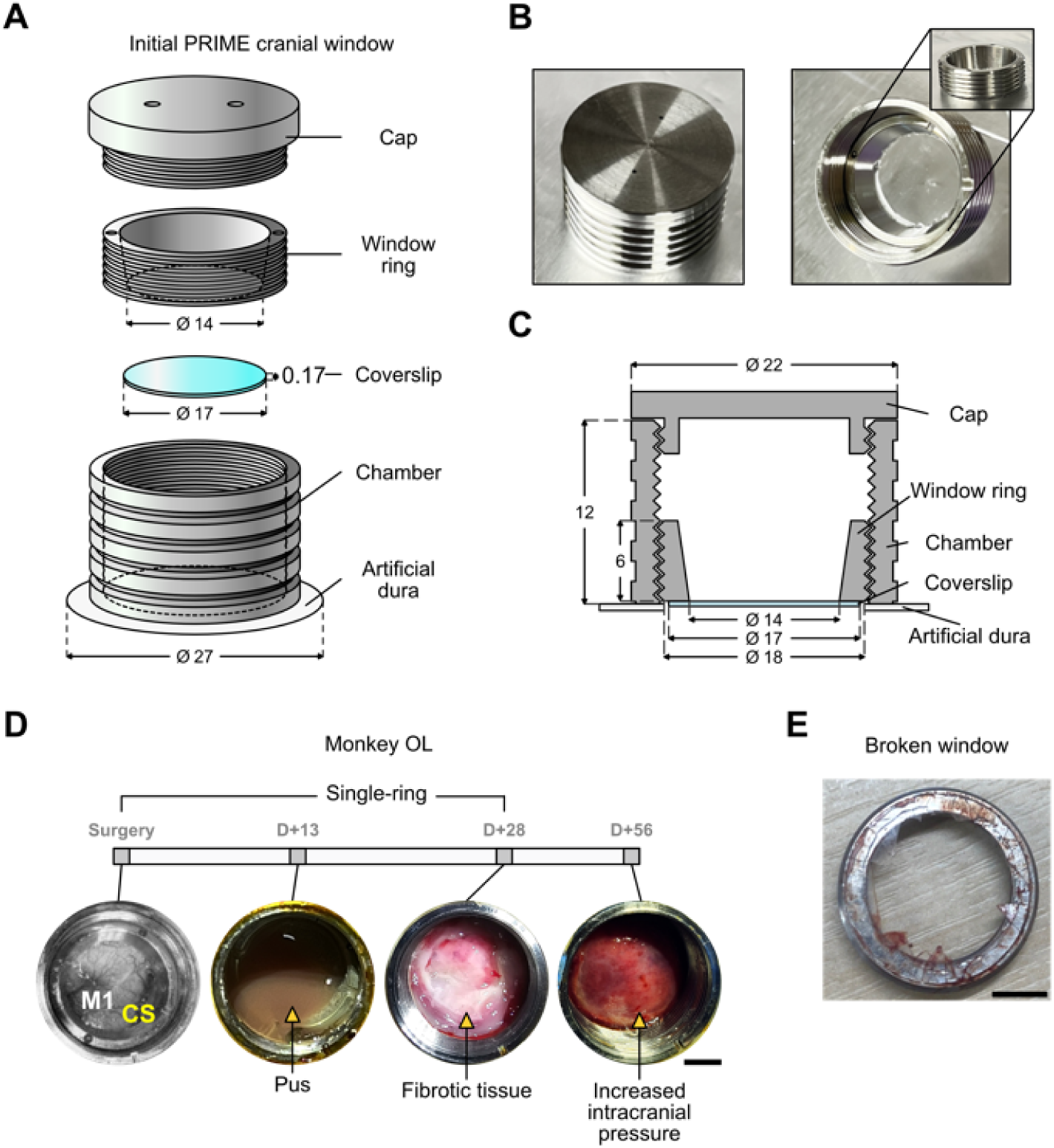
Single-ring cranial window and implantation outcome. Related to Figure 2. **A.** Initial single-ring cranial window design. A three-dimensional schematic shows the cap, window ring, coverslip, chamber, and artificial dura comprising the initial prototype. **B.** Fabricated prototype components. Representative images show the fabricated components of the initial single-ring cranial window. **C.** Single-ring window assembly. A cross-sectional schematic shows the arrangement and dimensions of the assembled prototype. **D.** Longitudinal implantation outcome. Serial monitoring in monkey OL shows progressive loss of cortical visibility associated with pus accumulation, fibrotic tissue formation, and increased intracranial pressure. **E.** Coverslip fracture. A representative image shows the fractured coverslip observed following the increase in intracranial pressure shown in (D). Scale bars, 5 mm. All dimensions are in millimeters.

**Figure S2.**
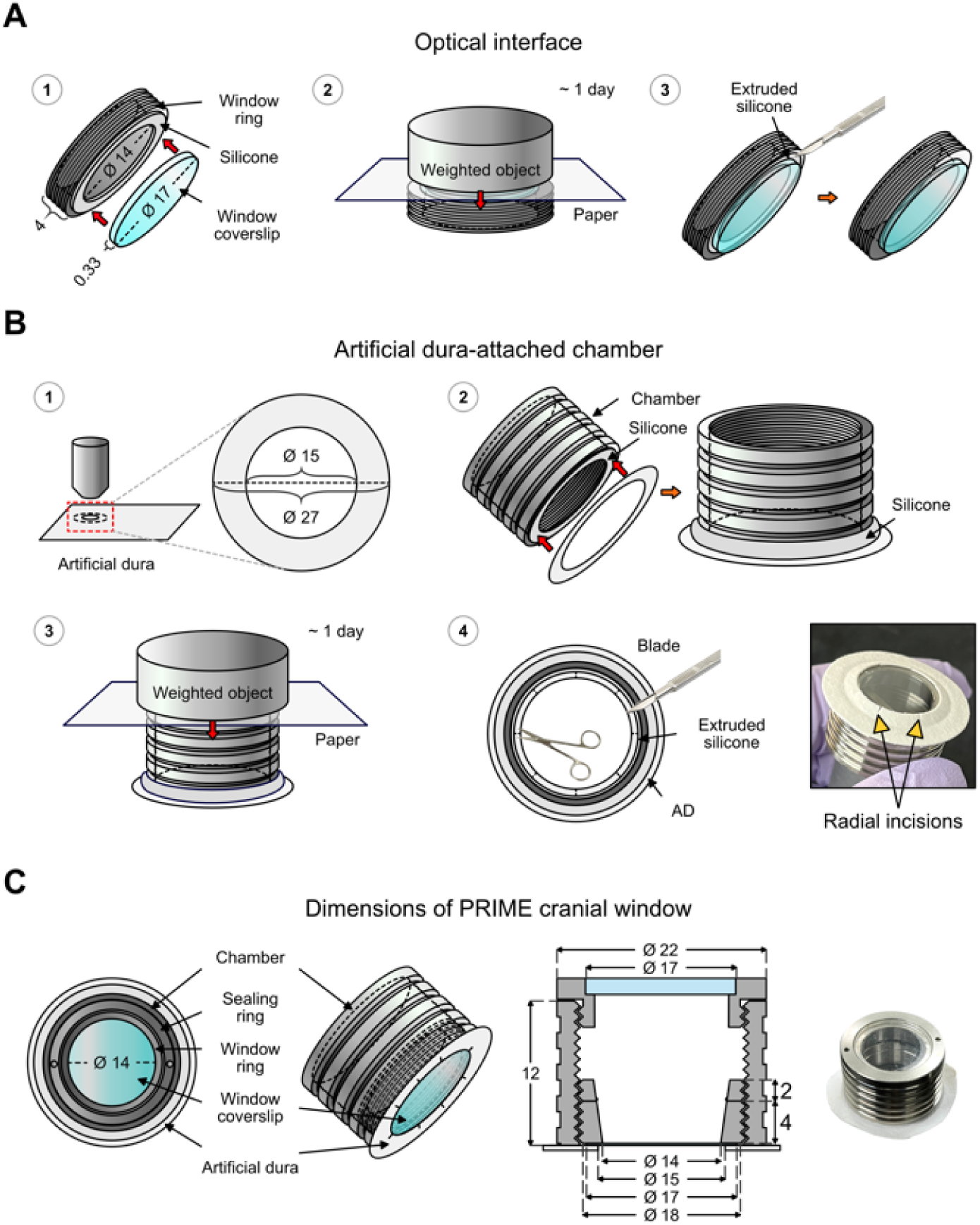
Fabrication and assembly of PRIME cranial window. Related to Figure 2. **A.** Optical interface fabrication. The coverslip was bonded to the window ring, compressed during adhesive curing, and trimmed to remove excess adhesive after curing. **B.** Artificial dura-attached chamber fabrication. The annular artificial dura was bonded beneath the chamber, compressed during adhesive curing, and trimmed before radial incisions were made along its inner margin. **C.** Final PRIME window dimensions. Schematics show the assembled optical interface, artificial dura-attached chamber, and final component dimensions, including a 14-mm optical aperture. All dimensions are in millimeters.

**Figure S3.**
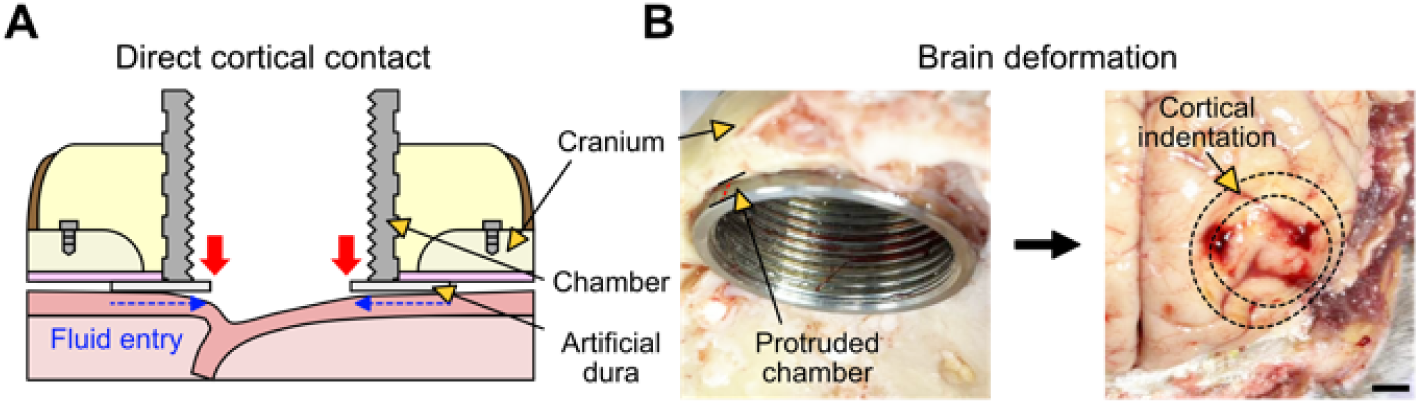
Brain deformation caused by chamber-cortex contact. Related to Figure 3. **A.** Direct chamber–cortex contact. Excessive chamber protrusion toward the brain produced focal indentation of the cortical surface. **B.** Altered brain–window apposition. Cortical deformation was accompanied by incomplete apposition of the artificial dura to the cortical surface and fluid entry beneath the window. Scale bar, 5 mm.

**Figure S4.**
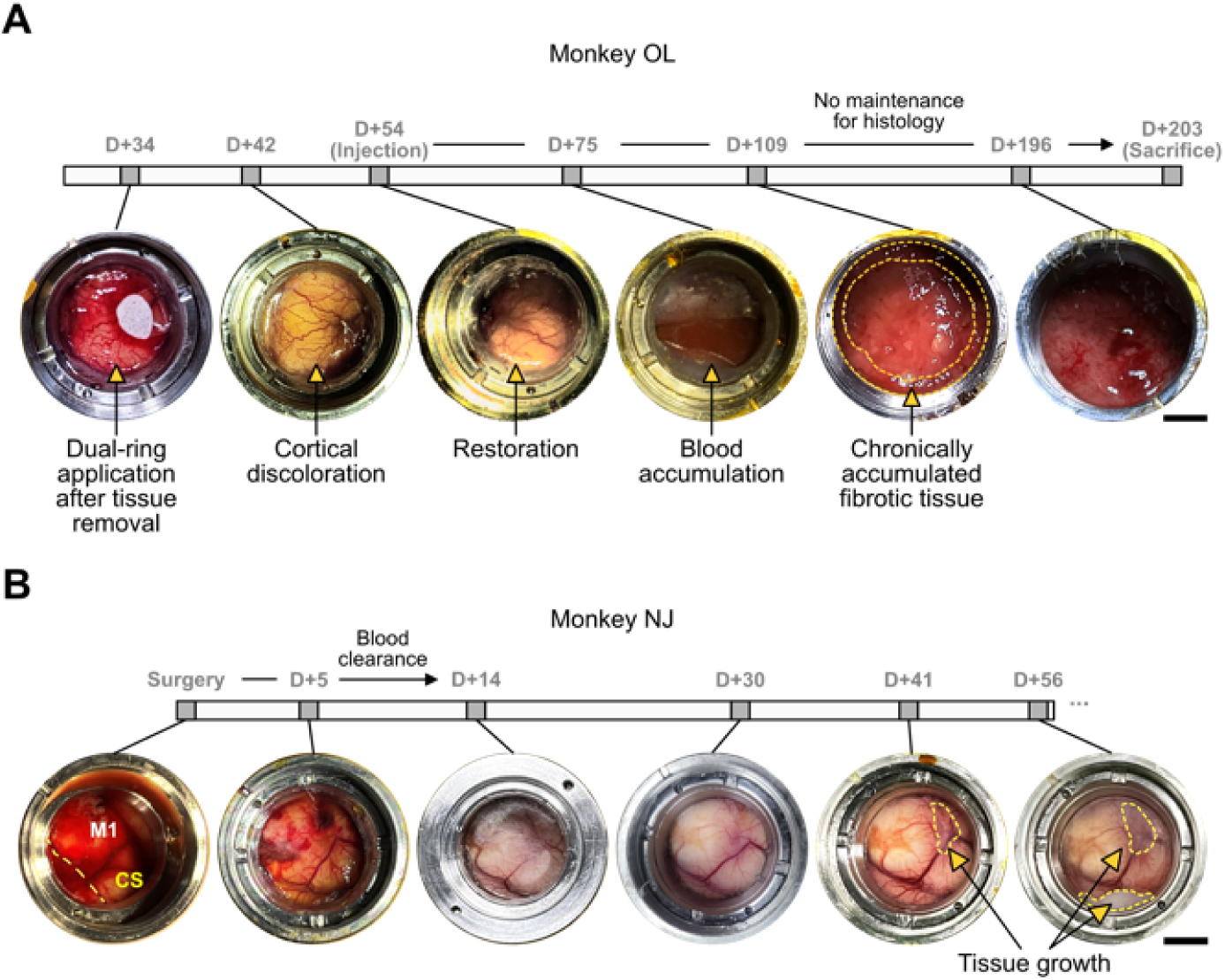
Longitudinal cortical responses after PRIME window implantation. Related to Figure 4. **A.** Longitudinal cortical response in monkey OL. Serial observations show cortical discoloration and recovery after tissue removal, followed by blood accumulation and chronic fibrotic tissue formation. Maintenance was withheld after post-implantation viral vector delivery to preserve the tissue condition for subsequent histological analysis. **B.** Longitudinal cortical response in monkey NJ. Serial observations show postoperative blood clearance followed by progressive tissue growth beneath the PRIME cranial window. CS, central sulcus; M1, primary motor cortex. Scale bars, 5 mm.

**Figure S5.**
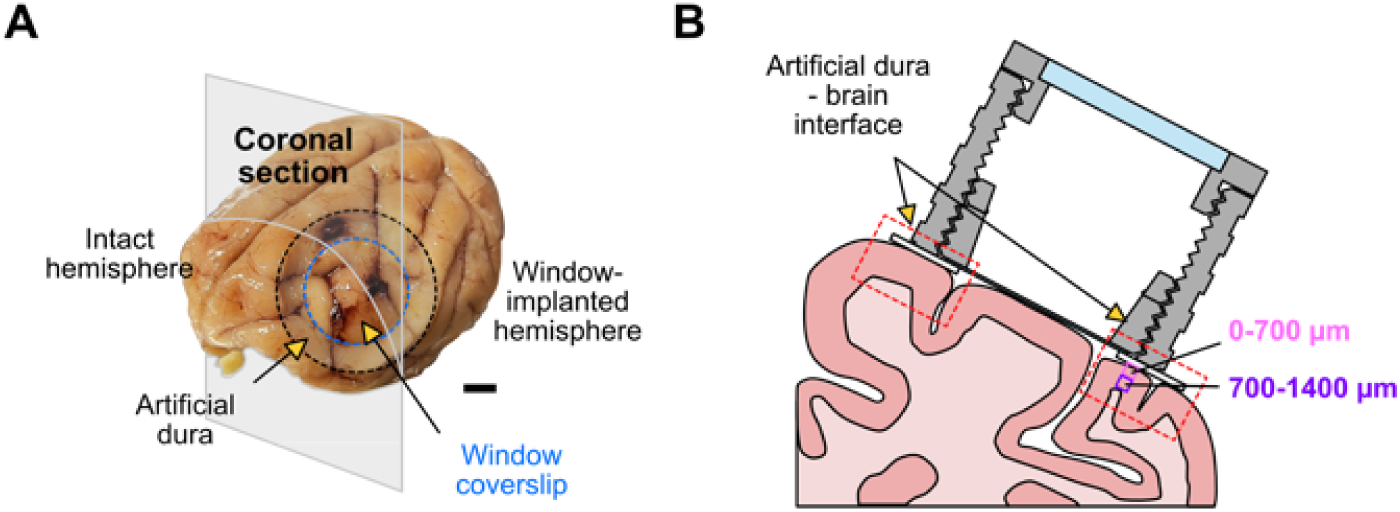
Histological ROI definition for artificial dura-associated immune-response analysis. Related to Figure 6. **A.** Histological comparison regions. A coronal brain section shows the hemisphere implanted with the PRIME window and the corresponding region in the contralateral intact hemisphere, with the artificial dura and window coverslip indicated. **B.** Depth-stratified cortical ROI sampling. Square ROIs were placed beneath the artificial dura-contacting cortex and in the corresponding intact cortex. Superficial and deeper cortical regions were defined as 0–700 μm and 700–1,400 μm from the pial surface, respectively. Scale bar in (A), 5 mm.

**Figure S6.**
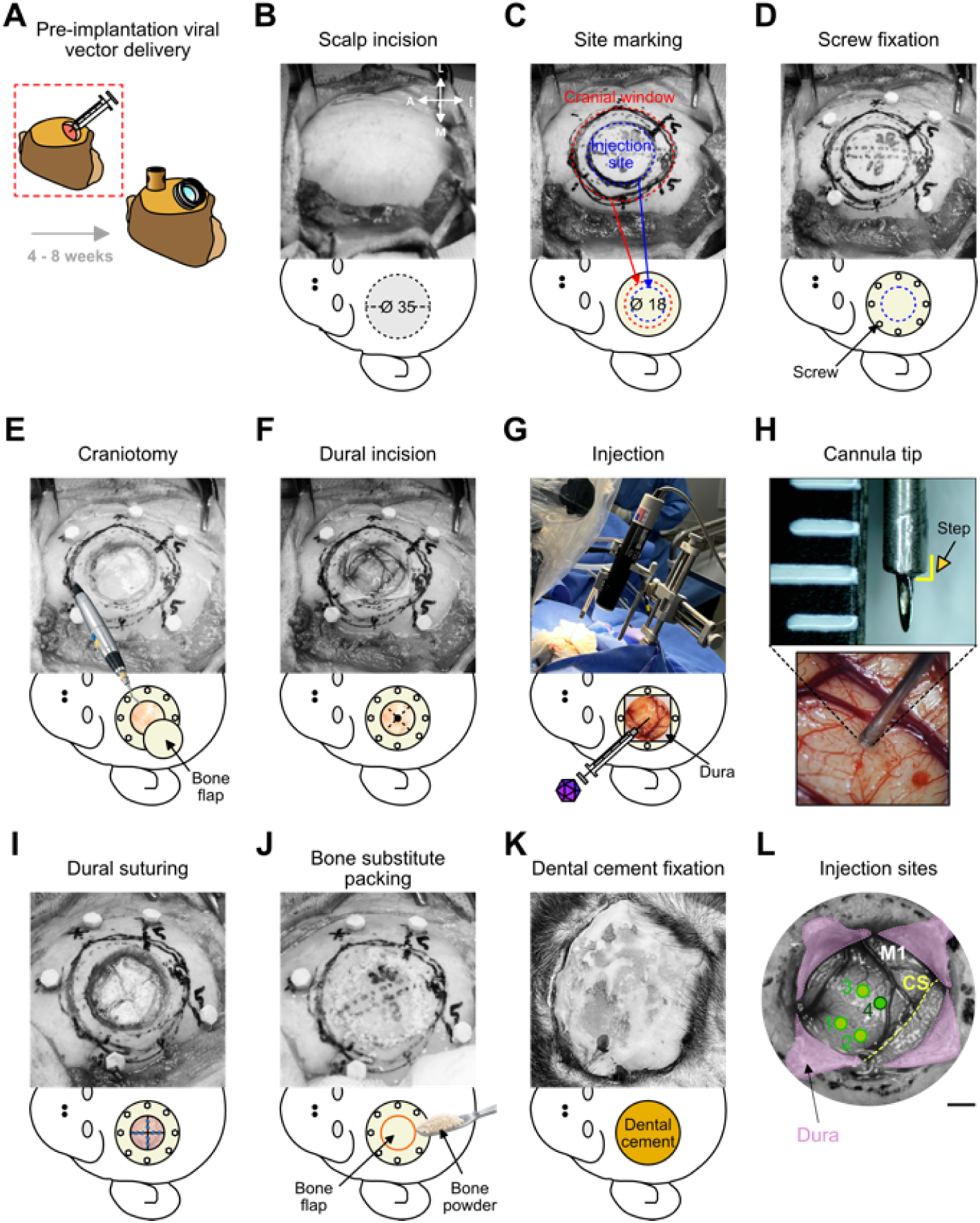
Schematic workflow for pre-implantation viral vector delivery. Related to Figure 7. **A.** Pre-implantation viral vector delivery strategy. Viral vectors were delivered 4–8 weeks before PRIME cranial window implantation to allow transgene expression before chronic optical access. **B.** Scalp incision and cranial exposure. The scalp was incised to expose the cranium over the planned injection and cranial window sites. **C.** MRI-guided site planning. The future cranial window boundary and injection region were marked over the target cortical area. **D.** Screw fixation. Fixation screws were positioned around the planned cranial window site before craniotomy. **E.** Bone-flap craniotomy. A craniotomy was performed at the injection site while preserving the bone flap for subsequent repositioning. **F.** Cruciate dural incision. The native dura was incised in a cruciate pattern while preserving the dural flaps for closure after injection. **G.** Stereotaxic viral vector injection. Viral vectors were delivered using a frame-mounted Hamilton syringe and injection pump. **H.** Stepped cannula insertion. Representative images show the stepped cannula tip and its insertion into the exposed cortical surface. **I.** Dural closure. The native dural flaps were repositioned and sutured after completion of the injections. **J.** Bone-flap replacement. The preserved bone flap was repositioned, and bone substitute was packed into the surrounding craniotomy gap. **K.** Dental cement fixation. Dental cement was applied over the reconstructed cranial site to complete closure. **L.** M1 injection map. Four viral vector injection sites were distributed across M1. Light green indicates AAV2.1-A-mCaMKIIα-ChRger2-mClover injection sites, and dark green indicates an AAV2/9-CaMKIIα-hChR2-eGFP injection site. All injections were performed at a depth of 1 mm. Site 1 received 3 μL at a rate of 1–5 μL min⁻¹, whereas sites 2–4 each received 1 μL at a rate of 0.2 μL min⁻¹. CS, central sulcus; M1, primary motor cortex. Scale bar in (L), 5 mm. All dimensions are in millimeters.

